# Profiling anti-apoptotic protein BCL-xL expression in U-87 MG glioblastoma cells and patient-derived tumorspheres

**DOI:** 10.1101/2020.03.24.005868

**Authors:** D. Fanfone, M. Gabut, G. Ichim

## Abstract

Glioblastoma is one of the cancers with the worst prognosis, despite huge efforts to understand its unusual heterogeneity and aggressiveness. These are mainly attributable to glioblastoma stem cells, which are also responsible for the frequent tumour recurrence following surgery or chemo/radiotherapy. We report here that tumorspheres derived from the U-87 MG glioblastoma cells have an increased expression of the anti-apoptotic protein BCL-xL. Modulation of this expression in tumorspheres enlightened us on the potential role of apoptosis and BCL-2 proteins might play in the survival of glioblastoma stem cells. Moreover, increased BCL-xL expression appears to sensitise glioblastoma cells to the newly developed BH3 mimetics, opening new therapeutic perspectives for treating glioblastoma patients.

## Introduction

Glioblastoma (GBM) is the most frequent primary malignant and invasive adult brain tumour, with an extremely devastating prognosis since the five-year survival rate does not exceed five per cent (1). The current therapy (STUPP treatment) combines surgical resection of the primary tumour, radiotherapy and chemotherapy with the alkylating agent temozolomide (TMZ) (2). However, despite these aggressive treatments, GBM invariably relapse, iterative surgical removal is rarely feasible and no consensual medical treatment is available leading to tumour progression and patient death (3). GBM recurrence can partially be explained by the important cellular complexity of GBM tumours and by a high inter-and intra-tumoral heterogeneity (4). Due to their pronounced cellular and molecular heterogeneity, GBM cells can adapt to selective pressures and changes in their microenvironment, leading to increased tumour aggressiveness and treatment resistance (5).

GBM tumours contain a small subset of cells with both stemness (self-renewing, proliferation, differentiation) and tumour-initiating properties, called glioblastoma stem cells (GSCs), driving tumour recurrence in GBM patients even after STUPP treatment (6) (7). Similarly to neural stem cells which have the capacity to form neurospheres *in vitro*, GSCs have the ability to form clonogenic structures, called tumorspheres, in serum-free non-adherent conditions *in vitro* (6) (8). These 3D culture models therefore represent models of choice to unravel and test new therapeutic strategies to specifically target GSCs and prevent GBM development (9).

Cancer cells must overcome several cellular stresses such as DNA damage, inflammation, oncogene activation, aberrant cell cycle progression and a harsh microenvironment that would normally trigger apoptosis in non-transformed cells (10). Apoptosis, the most extensively studied form of programmed cell death, is essential in maintaining the homeostatic balance between cell proliferation and cell death. Apoptosis can be triggered by two major signalling pathways, both leading to effector caspases activation and subsequent DNA fragmentation, plasma membrane blebbing and cell shrinkage (11). The extrinsic pathway is induced upon activation of death receptors (superfamily of tumour necrosis factor receptors) on the cell surface (12). The intrinsic pathway can be activated through various signals such as DNA damage, reactive oxygen species (ROS), or growth factor deprivation (11). Following mitochondrial outer-membrane permeabilization (MOMP), cytochrome *c* is thus released into the cytosol and engages the formation of the apoptosome that will first activate initiator caspase-9 and then effector caspases-3 and -7. These two pathways are controlled by BCL-2 family members: the pro-apoptotic (BAX, BAK, BID or BAD) and the anti-apoptotic proteins (BCL-2, BCL-xL, BCL-w, MCL-1) (11) (13) (14).

Cancer cells commonly share the ability to escape to programmed cell death. This evasion allows cancer cells to grow and develop into a tumour, while also contributing to treatment resistance (15). The blockade of cell death is a frequent cause of treatment resistance in GBM (16) (17) (18) (19).

Apoptosis escape commonly occurs through the deregulation of pro- and anti-apoptotic proteins in human cancers cells. The overexpression of anti-apoptotic proteins at transcriptional and protein levels has been observed in various cancers. BCL-2 was first described to be constitutively expressed in follicular lymphoma and the amplification of *MCL1* and *BCL2L1* (encoding BCL-xL) are the most frequent in solid cancers (13). In glioblastoma, MCL-1 is also overexpressed while high BCL-xL expression is often associated with poor prognosis and advanced disease (20) (21). Additionally, BCL-xL expression was also shown to increase with chemotherapy and ionizing radiation in lung cancer. Its role in stemness and aggressiveness is documented in melanoma and glioblastoma (22). Recently, BH3 mimetics developed to functionally imitate the pro-apoptotic BCL-2 proteins were shown to neutralize the anti-apoptotic proteins, enabling efficient apoptosis in cancer cells (23).

Considering its important function in regulating the apoptotic response in several cancers, we therefore focused on characterizing the expression and possible role of BCL-xL in GSCs growth and possible resistance to BH3 mimetics. The main finding of this short investigative work is that BCL-xL is highly expressed in tumorspheres originating from U-87 MG GBM cells, rendering them specifically sensitive to the BCL-xL inhibitor ABT-263. Although these findings are in our hands limited to one GBM cell line, this study provides interesting preliminary data for future research into repurposing BH3 mimetics for GBM treatment.

## Results

### High diversity of BCL-xL expression among GBM cell lines

As several research articles highlighted a link between resistance to apoptosis and cancer development in GBM, we speculated that anti-apoptotic proteins could be implicated in resistance to the therapy of CSC-like cells. Several protocols to isolate and culture GBM CSCs are described and currently used, however the most common one is culturing GBM cells in serum-free media, complemented by EGF and FGF which favours GBM tumorspheres formation (**Fig. 1 A**).

**Figure 1.**
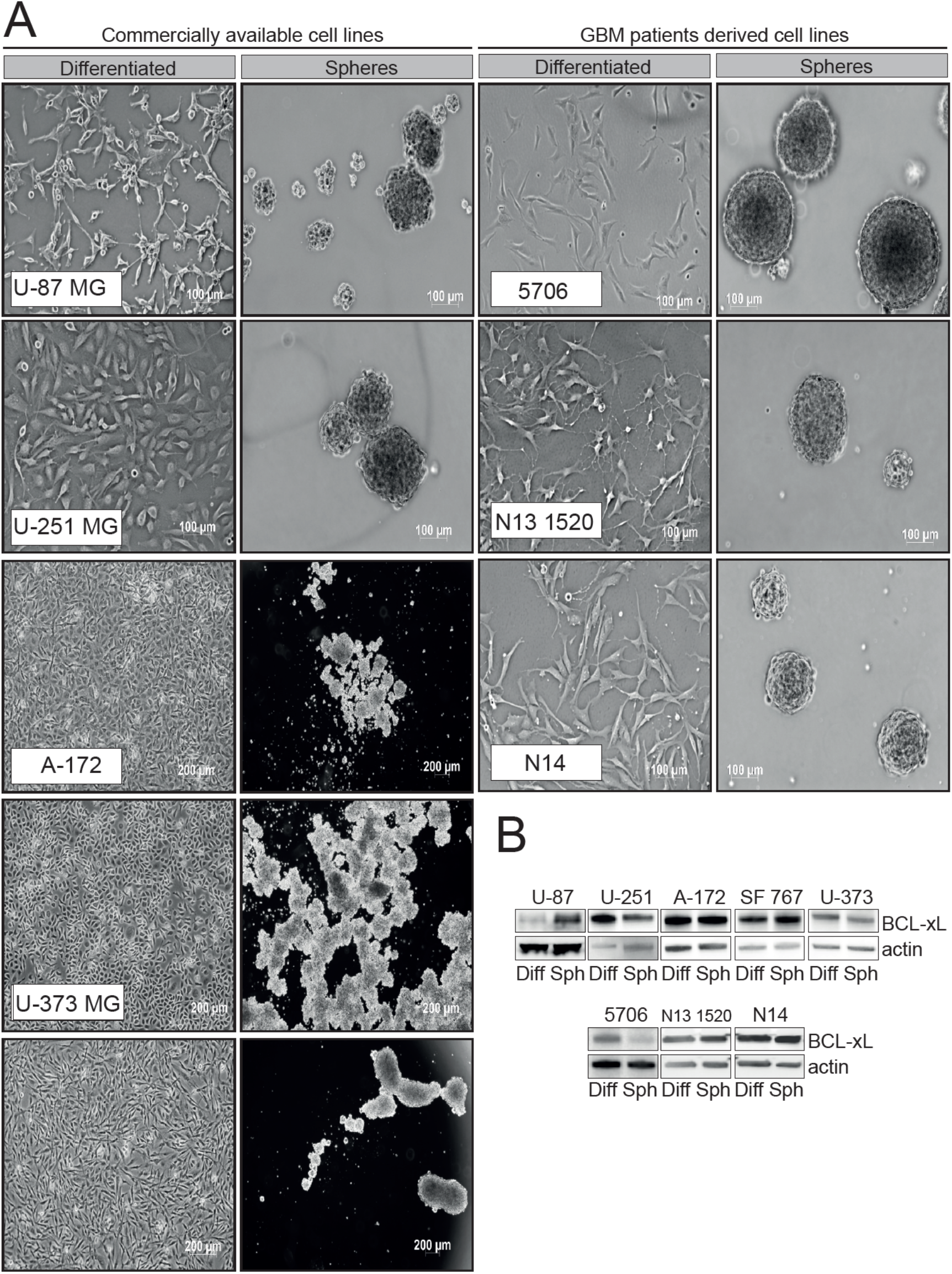
Evaluation of BCL-xL expression in various commercially available and GBM patients-derived cell lines. **A**. Images of GBM cell lines cultured as differentiated cells or their derived tumorspheres. **B**. Western blot analysis of BCL-xL expression in commercially available and GBM patients derived cell lines.

We sought to investigate BCL-xL expression in several GBM cell lines, either commercially available or patient-derived GSC tumorspheres cultures. These cells were either grown as adherent, differentiated cells, or allowed to form spheres when deprived of serum (**Fig. 1 A**). Given the widely accepted heterogeneity between GBM subtypes and even within the same tumour, it came as no surprise that all types of cells we tested show a different pattern of BCL-xL expression. This result highlights that the complexity of GBM is also mirrored in the different patterns of BCL-xL expression among various GBM cells. Of interest, the U-87 MG cells show a clear increase of BCL-xL expression when grown as tumorspheres (**Fig. 1 B**). We thus focused on this cell line as a model to investigate what might be the functions of BCL-xL in GBM tumorspheres.

### U-87 MG glioblastoma cells-derived spheres upregulate the anti-apoptotic protein BCL-xL

U-87 MG cells, widely used for *in vitro* GBM studies, were grown either in adherent standard conditions or in stem cell media (serum-free, supplemented with EGF and bFGF) and tumorspheres were collected after 7 and 14 days of culture. Since our initial aim was to assess the level of expression of anti-apoptotic protein in GBM tumorspheres, we analysed by western blotting the expression of the most important anti-apoptotic proteins, BCL-xL and MCL-1. Importantly, **Figure 2 A** shows a specific and significant upregulation of BCL-xL at the protein level in 7 and 14 days-old tumorspheres compared with differentiated U-87 MG cells, while MCL1 protein levels were not significantly changed. BCL-xL protein upregulation in tumorspheres is also mirrored at the mRNA level by an increase in *BCL2L1* mRNA, as assessed by qRT-PCR (**Fig. 2 B**). To exclude an indirect impact of EGF and bFGF added for tumorspheres culture, we tested the effect of these cytokines on the expression on BCL-xL, however no effect was noticed on its protein expression (**Fig. 2 C**).

**Figure 2:**
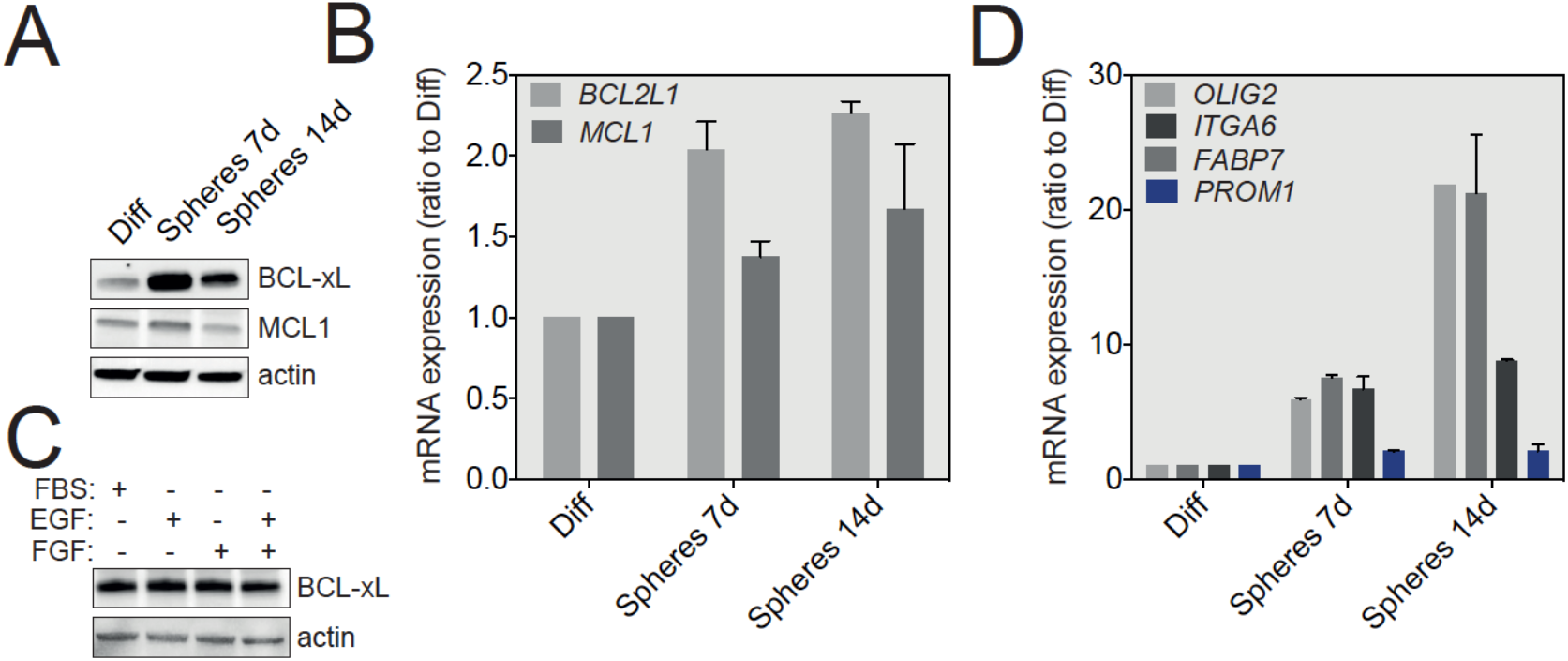
BCL-xL is highly expressed in U-87 MG-derived tumorspheres compared to differentiated cells. **A**. Western blot analysis of BCL-xL and MCL-1 expression in U-87 MG cells (differentiated versus tumorspheres). **B** Q-RT-PCR analysis of *BCL2L1* and *MCL1* in differentiated and spheres grown from U-87 MG cells at 7 and 14 days in culture. **C**. Western blot analysis of BCL-xL to test the influence of various growth factors used to grow tumorspheres. **D**. Q-RT-PCR analysis of selected stemness signature (*OLIG2, ITGA6, FABP7, PROM1*) in U-87 MG cells, grown as described in **B**.

We next assessed the levels of mRNA expression of several cell lineage specific proteins enriched in GSCs, including OLIG2, ITGA6, FABP7 and CD133 (*PROM1* gene) by qRT-PCR. As shown in **Figure 2 D**, U-87 MG-derived tumorspheres revealed an upregulation of all above-mentioned stemness markers that is directly correlated to the duration of tumorspheres culture, compared with adherent cells cultured in the presence of serum. These initial results therefore strongly suggest that U-87 MG tumorspheres support the growth of cells with GSC-like molecular signatures. Taken together, these results show that GSC-enriched U-87 MG-derived tumorspheres highly express the anti-apoptotic BCL-xL protein.

### BCL-xL regulates the size of U-87 MG neurospheres

We next investigated the possible roles that BCL-xL might play in the biology of GBM tumorspheres. For this, we first took advantage of U-87 MG cells stably expressing a degradation-sensitive DD-FLAG-BCL-xL that is therefore constantly degraded in the absence of Shield-1 (24). To finely tune the over-expression of BCL-xL in spheres, we first tested various concentrations of Shield-1 and choose 100 nM as working concentration to induce a significant BCL-xL protein accumulation for future experiments (**Figure 3 A**). Next, we treated U-87 MG derived tumorspheres with Shield-1 for 7 or 14 days (**Figure 3 B**) and then quantified tumorspheres size as indicator of their growth capacity. As depicted in **Figure 3 C** and confirmed by size quantification (tumorspheres average area) in **Figure 3 D-E**, BCL-xL overexpression significantly increases the size of U-87 MG tumorspheres. Intriguingly, the increased expression of BCL-xL does not affect the stemness markers (*OLIG2*, *ITGA6*, *FABP7* and *PROM1*) (**Fig. 3 F**). Since GBM spheres are composed of a mixture of GSCs and differentiated cells, we speculated that BCL-xL might impact on the proliferation of the latter cellular population. We next tested the effect of BCL-xL downregulation by stably expressing two lentiviral vectors expressing BCL-xL-targeting shRNAs. First, the efficacy of BCL-xL knockdown was validated both in differentiated cells and U-87 MG tumorspheres (**Fig. 4 A**). Of note, BCL-xL knockdown does not affect cell proliferation in the differentiated growth conditions (**Supp. Fig. 1 A**), while it clearly sensitize U-87 MG cells to actinomycin D induced apoptosis (**Supp. Fig. 1 B**). Actinomycin D is a chemotherapy drug blocking DNA transcription and therefore inducing apoptosis. Interestingly, knock-down of BCL-xL has the opposite effect on U-87 MG tumorsphere growth compared to its overexpression, namely the size of tumorspheres is significantly impaired (**Fig. 4 B, C**), again without a noticeable effect on the expression on stemness markers (**Fig. 4 D**). In summary, these results suggest that in a model of U-87 MG tumorsphere formation, BCL-xL plays a stemness-independent role in the modulation of GBM tumorspheres size.

**Figure 3:**
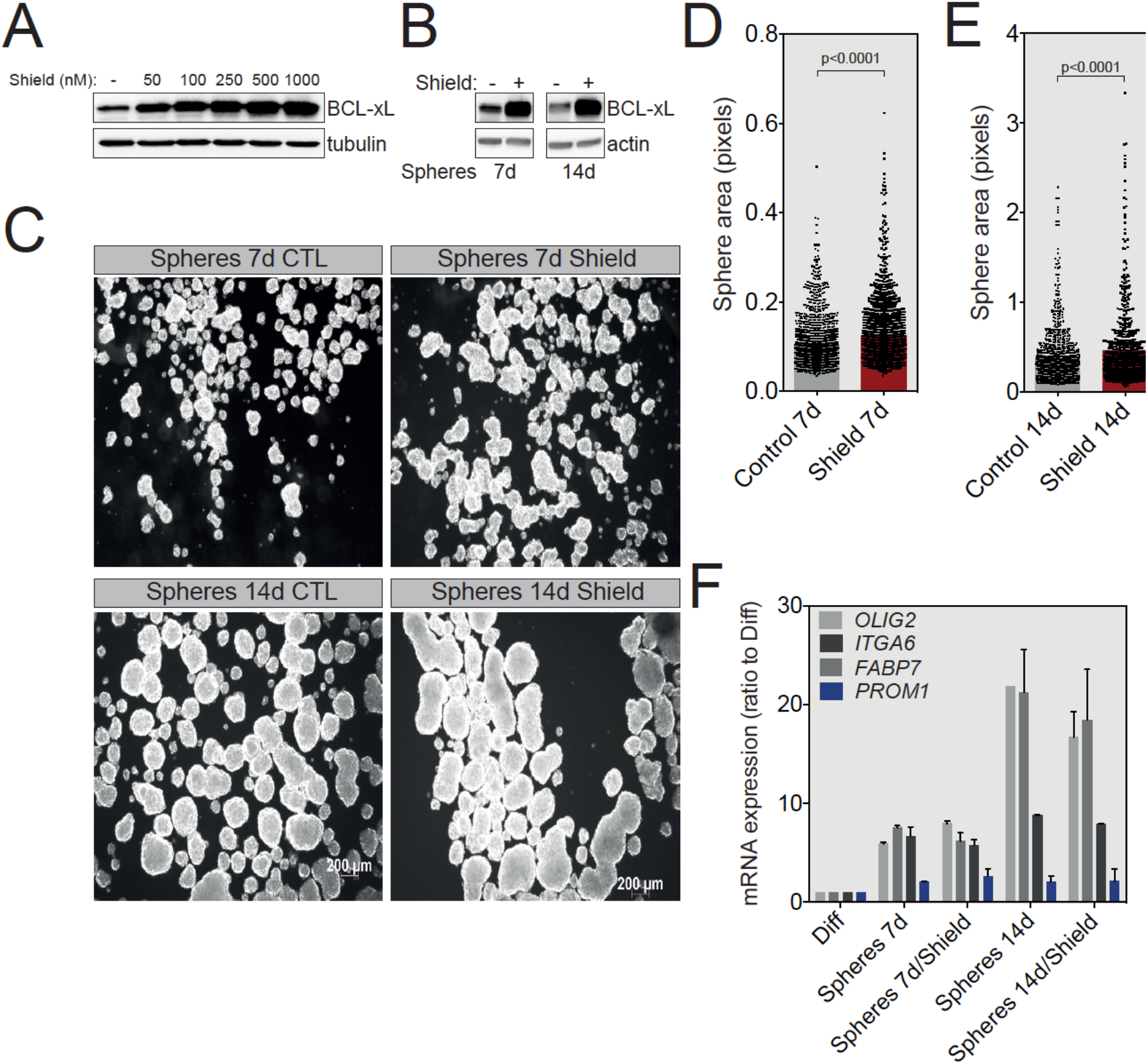
BCL-xL overexpression increases the number of U-87 MG-derived tumorspheres. **A**. Western blot analysis correlating the expression of BCL-xL with the concentration of Shield-1 ligand used to treat U-87 MG BCL-xLDD cells for 24 hours. **B**. Western blot analysis of BCL-xL overexpression in U-87 MG tumorspheres after 7 and 14 days of shield-1 treatment at the indicated concentration. **C**. Representative pictures of U-87 MG tumorspheres following 7 and 14 days of culture in the presence of shield-1. **D**. Comparison of the size of U-87 MG tumorspheres after 7 (**D**) or 14 (**E**) days of culture with and without Shield-1. **F**. Q-RT-PCR analysis of the stemness signature (*OLIG2, ITGA6, FABP7, PROM1*) in U-87 MG cells cultured in the absence or presence of Shield-1.

**Figure 4.**
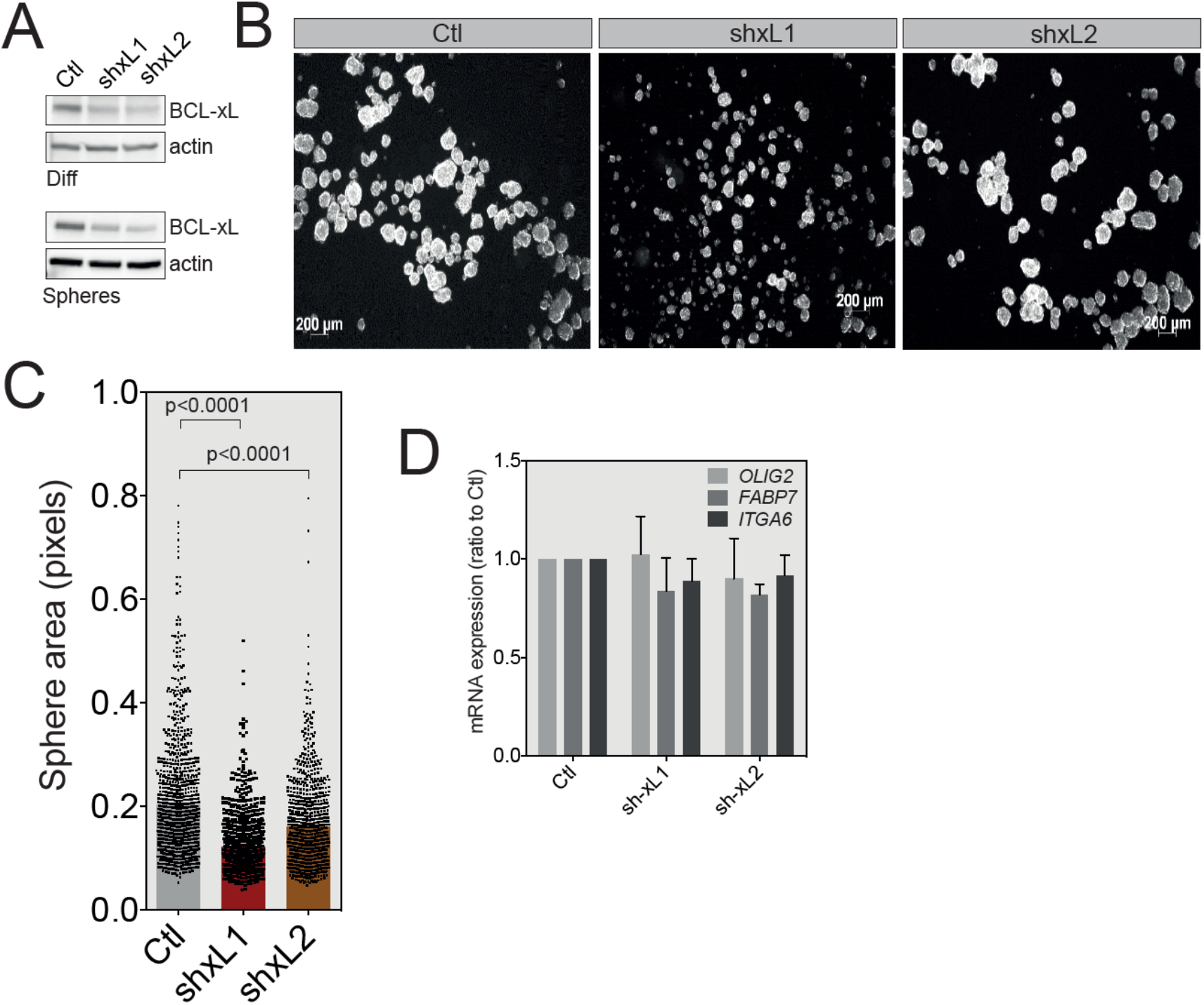
BCL-xL knock-down reduces GBM tumorspheres size. **A**. Western blot analysis of BCL-xL expression in differentiated or U-87 MG-derived tumorspheres following shRNA-mediated BCL-xL knock-down. Two different specific shRNA were employed and resulted in the same silencing efficacy. **B**. Representative images of U-87 MG tumorspheres after BCL-xL silencing. **C**. Comparison of the size of U-87 MG tumorspheres before and after BCL-xL knockdown. **D**. Q-RT-PCR analysis of the stemness signature (*OLIG2, FABP7, ITGA6*) in control cells versus BCL-xL knockdown tumorspheres.

### BCL-xL upregulation in GBM tumorspheres enhances the therapeutic targeting of GBM using BH3 mimetics

BH3 mimetics are a novel class of drugs tailored to mimic the function of the BH3-only proteins and therefore targeting the pro-survival members of the BCL-2 family (23). Several BH3 mimetics are available, each displaying a different level of affinity for a specific anti-apoptotic protein (**Fig. 5 A**). To test whether the increased expression of BCL-xL in U-87 MG tumorspheres underlies a newly acquired sensitization to a specific BH3 mimetic, we treated these spheres with the ABT-263, ABT-737 or S63845 BH3 mimetics and assessed the induction of cell death using the IncuCyte live cell imaging coupled with SYTOX Green staining of apoptotic cells. As shown in **Figure 5 B, C**, U-87 MG tumorspheres undergo apoptosis mush faster (as early as one hour) and at a much higher rate when treated with ABT-263 and ABT-737, two of the BH3 mimetics targeting BCL-xL and BCL2/BCL-xL/BCL-w, respectively. Of interest, the MCL-1 inhibitor, S63845, has a lower killing efficacy (**Fig. 5 B-C**). In addition, a short treatment of only 3 hours with ABT-263, but not S63845, can efficiently activate the effector caspase-3, as shown by the caspase-3 processing and PARP-1 cleavage (**Figure 5 D**). Of note, the pan-caspase inhibitor Q-VD-OPh blocks this processing, therefore confirming an induction of apoptosis in tumorspheres treated with the BCL-xL inhibitor ABT-263. We have also performed a long-term tumorsphere survival assay, where U-87 MG spheres were briefly treated for 3 hours with either ABT-263 or S63854 and then allowed to grow in fresh media for 7 days. As depicted in the representative spheres pictures in and corresponding quantifications, the spheres obtained following ABT-263 treatment were both smaller in size (**Figure 5 E)**and fewer in number (**Figure 5 F**) than both control spheres and spheres treated with the MCL-1 inhibitor, indicating the long-term effect of this BH3 mimetic. Moreover, in this long-term survival assay, Q-VD-OPh did not rescue tumorsphere growth, exposing mitochondrial outer membrane permeabilization as the point of no return for tumorsphere cells apoptosis. To assess whether ABT-263 treatment affected the GSCs population, we conducted a qRT-PCR assay on tumorspheres treated with ABT-263 and observed that while there was no change for *ITGA6* and *FABP7*, expression of *OLIG2* increased sharply (**Fig. 5 G**). Collectively, these results provide important insights into targeting GBM spheres using tailored BH3 mimetics, owing to their elevated anti-apoptotic expression.

**Figure 5.**
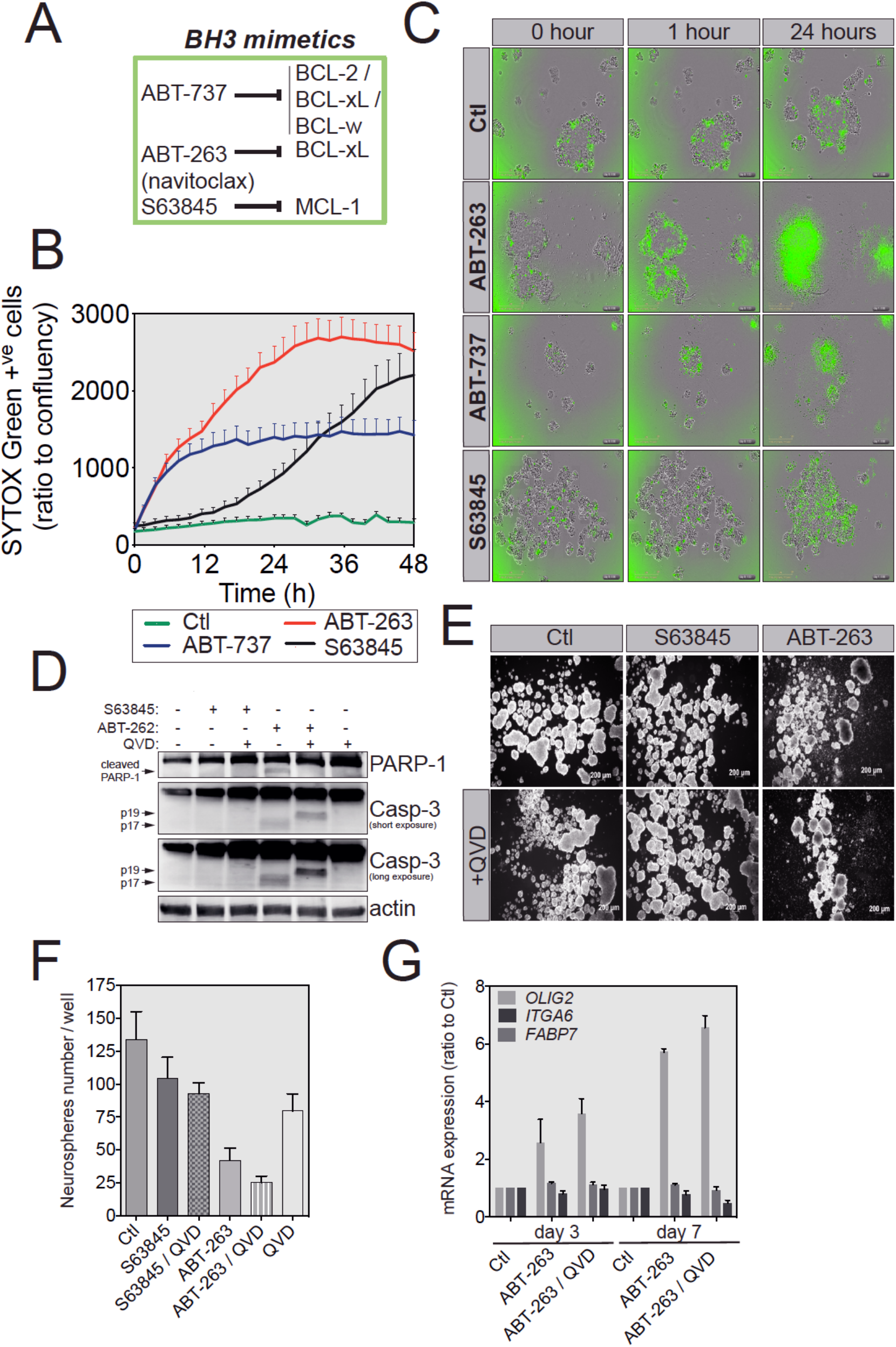
U-87 MG tumorspheres have an increased sensitivity to ABT-263-induced apoptosis. **A**. Brief summary of the main BH3-mimetics and their preferential targets. **B**. IncuCyte imager-based cell death induction analysis of U-87 MG-derived tumorspheres treated with ABT-737, ABT-263 or S63845 as indicated. Induction of apoptosis was assessed by SYTOX Green incorporation into permeabilized dead cells. **C**. Representative pictures of U-87 MG tumorspheres treated with different BH3-mimetics, while the green signal indicates SYTOX Green positive apoptotic cells. **D**. Western blot analysis of PARP-1 and caspase-3 protein expression and cleavage following treatment with BH3 mimetics. **E**. Representative images of U-87 MG tumorspheres in a long-term survival. Briefly, following treatment with the indicated BH3-mimetics, U-87 MG-derived tumorspheres were cultured in fresh medium for another week and imaged. **F**. Quantification of tumorspheres number for the long-term survival assay described in **E**. **G**. Q-RT-PCR analysis of stemness signature of U-87 MG tumorspheres treated with ATB263, in the absence or presence of 20 μM of pan-caspase inhibitor QVD-OPh.

## Discussion

In this report, we provide an investigative approach into the pattern of BCL-xL expression and possible function in GBM neurospheres, especially in U-87 MG cells. Compared to differentiated cells, U-87 MG tumorspheres express high levels of BCL-xL at both transcript and protein levels, independently of medium composition, suggesting an important role for BCL-xL in tumorspheres formation. Indeed, by artificially increasing BCL-xL expression in U-87 MG tumorspheres, we observed a significant increase in their size. Conversely, this was reverted when BCL-xL was downregulated *via* shRNA, and these latter cells were extremely sensitive to actinomycin D, which is highly reminiscent of results obtained with MCL-1 knockdown glioblastoma cells exposed to BH3 mimetics (20).

As confirmed by the upregulation of stemness genes, the population of GSCs is clearly more abundant in U-87 MG tumorspheres compared to adherent cells. However, we cannot yet prove at this stage that BCL-xL is specifically upregulated only in the GSCs compartment, given that tumorspheres are certainly composed of a mix of GSCs, progenitors and differentiated cells (25). To address this issue in future experiments, we would need to isolate from tumorspheres the two populations (differentiated and GSCs) and compare their respective expression of BCL-xL. This could be achieved for instance *via* flow cytometry-based cell sorting according to the expression of a cell surface marker specifically expressed by GSCs. However, although commonly described as a marker of GSCs, CD133 is probably not the most appropriate stemness marker in our hands and accordingly to other studies since only the glycosylated cell-surface CD133 epitope is stem cells-specific, while mRNA levels appear to be unrelated to stemness (26) (27) (28). However, OLIG2 and ITGA6 appear to be promising alternatives to faithfully characterize U87-MG GSCs, with ITGA6 being a possible plasma membrane marker that could allow cell sorting of GBM stem cells (29). In line with our findings in U87-MG cells, Trisciuglio and colleagues observed an increase in the amounts of tumorspheres number when BCL-xL is overexpressed in glioblastoma cells while the opposite was noticed when using BCL-xL depleted cells. They also noted that BCL-xL overexpression positively regulated the cancer stem cell phenotype in tumorspheres compared to control cells (22). These authors took their efforts further by demonstrating that exogenous BCL-xL controls several hallmarks of cancer aggressiveness such as migration, invasion and tumour cell plasticity, both in GBM and melanoma (22).

To improve our understanding of the transcriptional and/or translational regulation of BCL-xL in U87-MG neurospheres, an extensive study of various signalling pathway should be envisaged. Since BCL-xL can be regulated by a plethora of transcription factors ranging from STAT, Rel/NFkB or Ets, further insight into the mode of action of these regulators may unravel the specific regulation of BCL-xL in GBM tumorspheres (30).

A growing body of evidence suggests apoptosis-independent functions for several BCL2 family members, ranging from cell cycle, DNA damage response, metabolism or autophagy (31). In this article we focused on the anti-apoptotic function of BCL-xL and more specifically on revealing a therapeutic sensitivity to BH3 mimetics targeting BCL-xL. Indeed, the upregulation of BCL-xL in U-87 MG tumorspheres is linked to a higher sensitivity to BCL-xL specific BH3 mimetics. This finding corroborates a study by Liwak and colleagues that reported higher sensitivity of GBM cells to a combination of ABT-737 and doxorubicin, owing to increased levels of BCL-xL in a specific subset of glioblastoma cell lines (32). This is a highly promising finding, as it highlights means to kill GSCs, responsible for GBM tumours recurrence. However, these results should be interpreted with great caution as our findings are neither applicable to all patient-derived nor commercially available GBM cell line. BCL-xL can be targeted only if fresh tumour samples grown in tumorspheres permissive medium exhibit higher BCl-xL expression. Nevertheless, the use of BH3 mimetics for the treatment of glioblastoma could alleviate other pro-oncogenic effects of BCL-xL. For instance, BCL-xL was shown to function as an epigenetic modifier and promote both epithelial to mesenchymal transition (EMT) and stemness in pancreatic, breast and lung cancer (33) (34). This is also consistent with a growing number of pre-clinical studies confirming the benefits of using BH3-mimetics when treating gliomas (35).

## Materials and methods

### Cell lines and growth conditions

The U-87 MG, U-251 MG, U-373 MG, A-172, SF767 human glioblastoma cell lines were obtained from American Type Culture Collection (ATCC). The N13 1520, N14, 5706 are patients derived cell lines obtained from M. Gabut’s laboratory (CRCL). The human glioma cell lines were cultured in DMEM containing 2mM L-glutamine (ThermoFisher Scientific, 25030-24), non-essential amino acids (ThermoFisher Scientific, 11140-035), 1mM sodium pyruvate (ThermoFisher Scientific, 11360-039), 10% FBS (Eurobio, CVFSVF00-01) and 5% penicillin/streptomycin (ThermoFisher Scientific, 15140-122). Tumorspheres were obtained by culturing cells in stem cell medium (SCM) composed of DMEM F12 (ThermoFisher Scientific, 31331-093), B-27™ supplement 1x (Life Technologies, 17504044), EGF 20 ng/mL, recombinant human basic FGF 10 ng/mL (Peprotech, 100-18B) and 10% penicillin/streptomycin. Cells formed neurospheres only in plates with ultra-low attachment surface. In order to work in clonal conditions, 5×10^3^ cells were grown in 2 mL of SCM medium in a 6-wells plate. Cells were regularly checked for mycoplasma contamination.

### Reagents

Q-VD-OPh (Clinisciences, JM-1170), SYTOX Green (ThermoFisher Scientific, S34860), Actinomycin D (Sigma, A9415), ABT737 (Clinisciences, A8193), ABT263 (Navitoclax), S63845 (Clinisciences, A8737) were employed in this study.

### Stable cell lines generation

The retroviral constructs (pLZRS BCL-xL DD and pSuper.retro-shRNA BCL-xL were transfected into Amphotropic Phoenix cells (0.5 × 10^6^ in a 10 cm dish) using Lipofectamine 2000 (Invitrogen). Two days later virus-containing supernatant was harvested, filtered and used to infect target U-87 MG cells in the presence of polybrene (1 μg/mL). Two days post-infection, stably expressing cells were selected using Zeocin (200 μg/mL, Invitrogen). The sequence of shRNA BCL-xL 1 (or shxL1) is CCAGGAGAACCACTACATGCAGCC while that of shRNA BCL-xL 2 (or shxL2) is GTTCCAGCTCTTTGAAATAGTCTGT.

### Western blot analysis

Cell pellets were lysed in RIPA lysis buffer (Cell Signaling, 9806S) supplemented with phosphatase inhibitors complex 2 and 3 (Sigma Aldrich, P5726-1ML, P6044-1ML), DTT 10 mM and protease inhibitor cocktail (Sigma-Aldrich, 4693116001) in order to extract protein content. The quantification of proteins was performed using the Protein Assay dye Reagent Concentrate (Biorad, 50000006). Equal amounts (20 μg) of each samples were separated on 4-12% SDS-polyacrylamide gels (Biorad) under denaturating conditions (SDS PAGE Sample loading buffer (VWR, GENO786-701) supplemented with 1 mM DTT) and transferred onto a nitrocellulose membrane using the Transblot Turbo Transfer System (Biorad, 1704150EDU). Non-specific binding sites were blocked for 1 hour with 5% dry milk or BSA in TBS-Tween 0.1% while the primary antibody (1/1000 in 1% BSA TBS-Tween 0.1%) was incubated with the membranes overnight at 4°C, under agitation. The primary antibodies used were: actin (Sigma-Aldrich, A3854), BCL-xL (Cell Signaling, 2764S), MCL-1 (Cell Signaling, 4572S), beta-tubulin, PARP-1 (Cell Signaling, 9532), Caspase-3 (Cell Signaling, 9662S). Membranes were rinsed 3×10 minutes in TBST 0.1% then incubated with appropriate secondary antibody coupled to the horseradish peroxidase (Biorad, 1706515, 1706516; 1/5000) for 1 hour at room temperature under agitation. Three extra washing steps were performed before detection by chemiluminescence (Clarity Western ECL reagent, Biorad, 1705060) and chemiDoc Imager (Biorad, 17001401).

### Quantitative RT-PCR

Total RNA from cell pellets was extracted and purified using the Nucleospin RNA protocol from Macherey-Nagel kit (740955). Dosage of RNA samples was performed using a NanoDrop. mRNAs were converted to cDNA using the Sensifast cDNA synthesis kit (Bioline, BIO-65053). The cDNAs were then amplified by PCR by using specific primers for each gene designed on the Primer-blast software (https://www.ncbi.nlm.nih.gov/tools/primer-blast/) and listed in Table 1. *GAPDH* and *ACTB* were used as housekeeping genes. The PCR amplification was performed through an initial polymerase activation step at 95°C for 2 minutes, followed by 40 cycles at 95°C for 5 seconds and 60°C for 30 seconds. The q-RT-PCR experiments were conducted using SYBR Green and a Lightcycler96 (Roche, Indianapolis, USA).

**Table 1:**
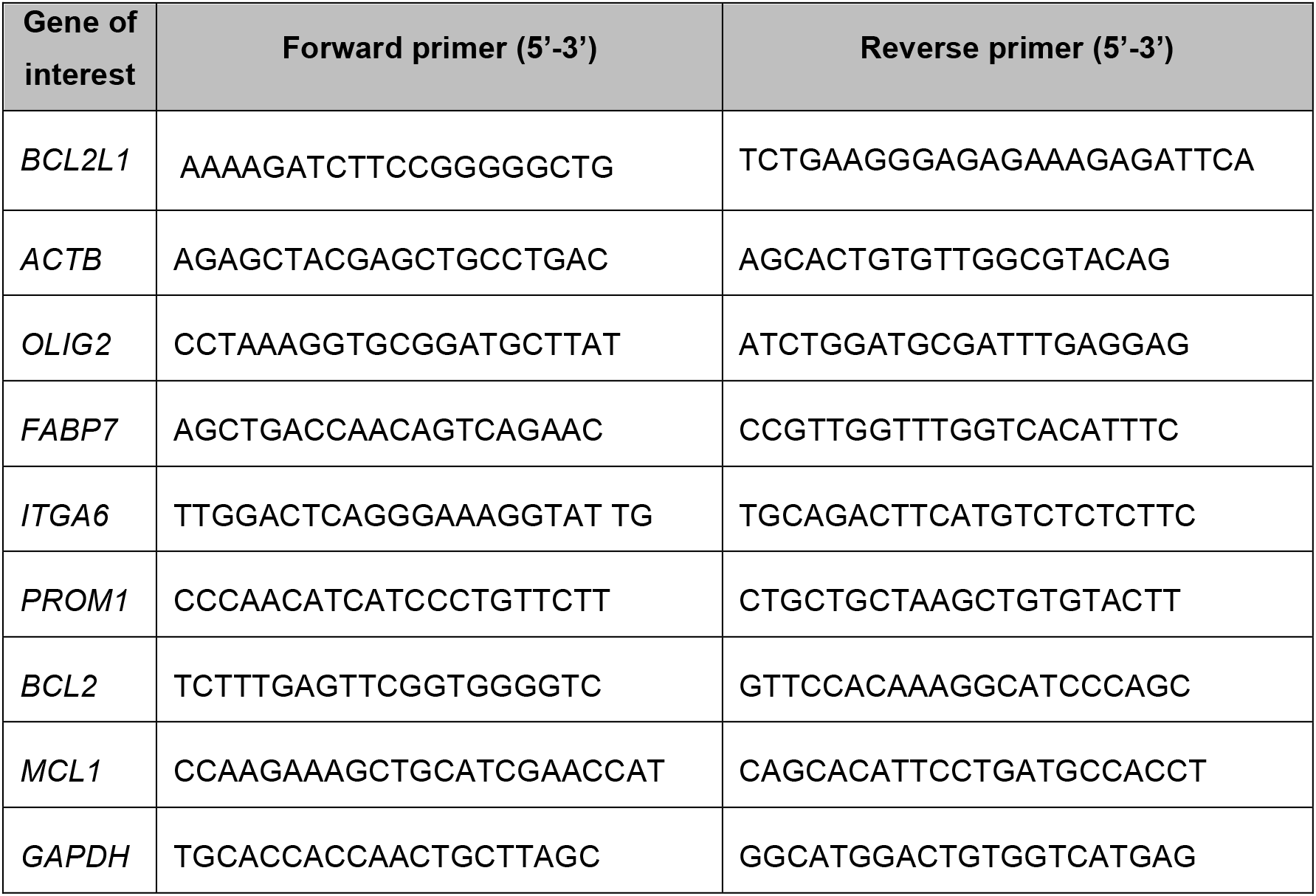
List of specific primers used for RT-qPCR.

### Apoptosis assay

Under a binocular microscope and in sterile conditions, 1-2 tumorspheres were placed in each well of a ULA 96-well plate (Greiner) and then treated with various compounds: 10 μM of ABT-737, ABT-263 or S63845, QVD-OPh 20 μM or with actinomycin D 1 μM in the presence of 30 nM SYTOX Green. Cells were then imaged every 60 minutes using the IncuCyte ZOOM imager.

### Image analysis

Measurements of tumorspheres size were performed using the ImageJ software 1.52a.

### Statistical analysis

Data are expressed as the mean ± SD. A two-tailed Student’s t-test was applied to compare two groups of data. Analyses were performed using the Prism 5.0 software (GraphPad).

---

**Supplementary Figure1 (related to Figure 3).**
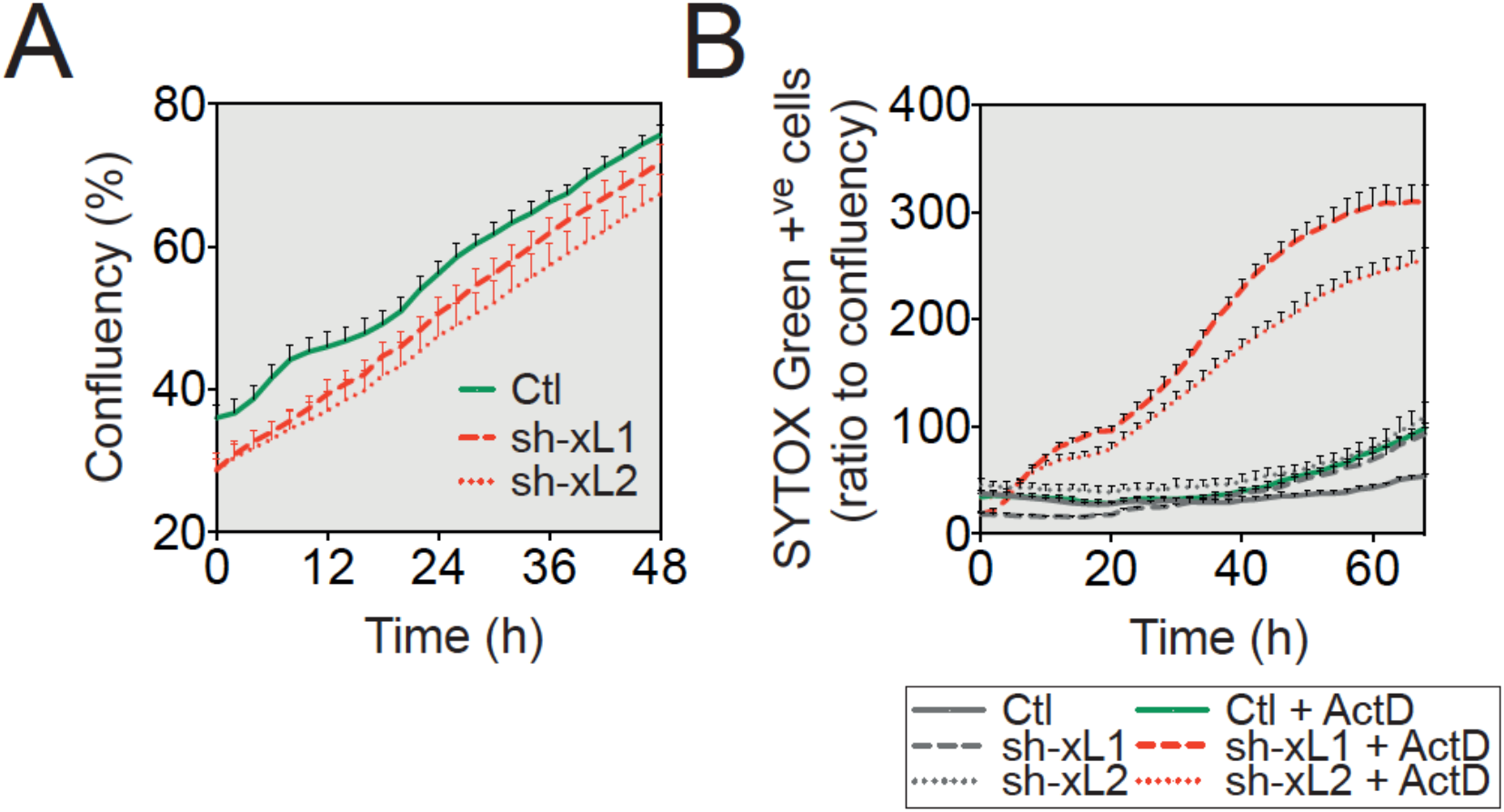
**A.** Cell growth analysis based on IncuCyte live imaging of control U-87 MG and shRNA BCL-xL cells. **B**. Apoptosis induction based on SYTOX Green incorporation following actinomycin D (1 μM) treatment.

## Acknowledgements

The authors would like to thank David Neves for providing cell lines, Germain Gillet and Nikolay Popgeorgiev for the shRNA BCL-xL plasmids, Stephen Tait for the DD-FLAG-BCL-xL plasmid, Nabila Berabez for technical assistance and Brigitte Manship for editing this article.

